# Kynurenine Metabolism is Associated with Antidepressant Response to Selective Serotonin Reuptake Inhibitors

**DOI:** 10.1101/2025.01.11.632543

**Authors:** Ryan Rampersaud, Klara Suneson, Gwyneth W. Y. Wu, Victor I Reus, Daniel Lindqvist, Tiffany C. Ho, Dieter J. Meyerhoff, Michael R. Irwin, Owen M. Wolkowitz, Synthia H. Mellon, Lena Brundin

**Author notes:** Corresponding author; Contact: Ryan Rampersaud MD PhD.

## Abstract

Alterations in the kynurenine pathway, and in particular the balance of neuroprotective and neurotoxic metabolites, have been implicated in the pathophysiology of Major Depressive Disorder (MDD) and antidepressant treatment response. In this study, we examined the relationship between changes in kynurenine pathway activity (Kynurenine/Tryptophan ratio), focusing on the balance of neuroprotective-to neurotoxic metabolites (Kynurenic Acid/Quinolinic Acid and Kynurenic Acid/3-Hydroxykynurenine ratios), and response to 8 weeks of selective serotonin reuptake inhibitor (SSRI) treatment, including early changes four weeks after SSRI initiation. Additionally, we examined relationships between kynurenine metabolite ratios and three promising biomarkers of depression and antidepressant response: amygdala/hippocampal volume, and glutamate metabolites in the anterior cingulate cortex. Responders showed an increase in the Kynurenic Acid/3-Hydroxykynurenine ratio by week 8 (*F*_(1,46)_ = 11.92, p = .001) and early increases in the Kynurenine/Tryptophan ratios at week 4 (*F*_(2,58)_ = 5.224, p = .008), while Non-Responders did not. Pre-treatment Kynurenic Acid/Quinolinic Acid and Kynurenic Acid/3-Hydroxykynurenine ratios were positively associated with right amygdala volume (β = . 247 p = .032 and β = .245 p = .028, respectively). Lastly, in a subset of participants, pre-treatment Kynurenic Acid/3-Hydroxykynurenine ratio showed a positive, small effect size association with glutamate metabolites (Glx) in the anterior cingulate cortex (β = .307 p = .079), which became significant post-treatment with a large effect size (β = .652 p = .021). These results suggest that response to SSRIs may arise from shifting the balance from neurotoxic to neuroprotective kynurenine metabolites.

## Introduction

Despite recent advances, a clear picture of the pathophysiology of depression has not yet emerged[1]. Although commonly prescribed[2], antidepressants fail to fully reduce the burden of disease with only 30 – 40% of individuals achieving remission with their first selective serotonin reuptake inhibitor (SSRI) trial[3,4] and diminishing likelihood of response with each successive trial[5]. Altered tryptophan metabolism, particularly alterations in the kynurenine pathway (**KP**), represents a novel pathophysiological mechanism in MDD and antidepressant treatment response[6] . These bioactive metabolites may serve as both markers guiding treatment selection as well as potential therapeutic targets.

Tryptophan, an essential amino acid, is primarily metabolized down the kynurenine pathway (>95%), with serotonin production representing a minor pathway[7] . The KP generates several neuroprotective metabolites (including Kynurenic Acid [KA] and Picolinic Acid [PA] as well as neurotoxic metabolites including 3-Hydroxykynurenine [3-HK], and Quinolinic Acid [QA]; **Figure 1**), and is preferentially shifted towards the neurotoxic branch of the KP by inflammatory mediators[8,9]. Rather than absolute abundance of KP-metabolites, the balance between the neuroprotective and neurotoxic arms of the KP may be of importance in the context of brain-based disorders, including depression[10–13]. For example, KA can act as a free radical scavenger, exert anti-inflammatory effects, or act as a NMDA receptor antagonist to protect against glutamate induced excitotoxicity[14]. In contrast, metabolites of the neurotoxic branch can stimulate production of free radicals, induce mitochondrial dysfunction and apoptosis, promote inflammation, or act as an NMDA receptor agonist potentially contributing to glutamate induced excitotoxic damage[15].

**Figure 1.**
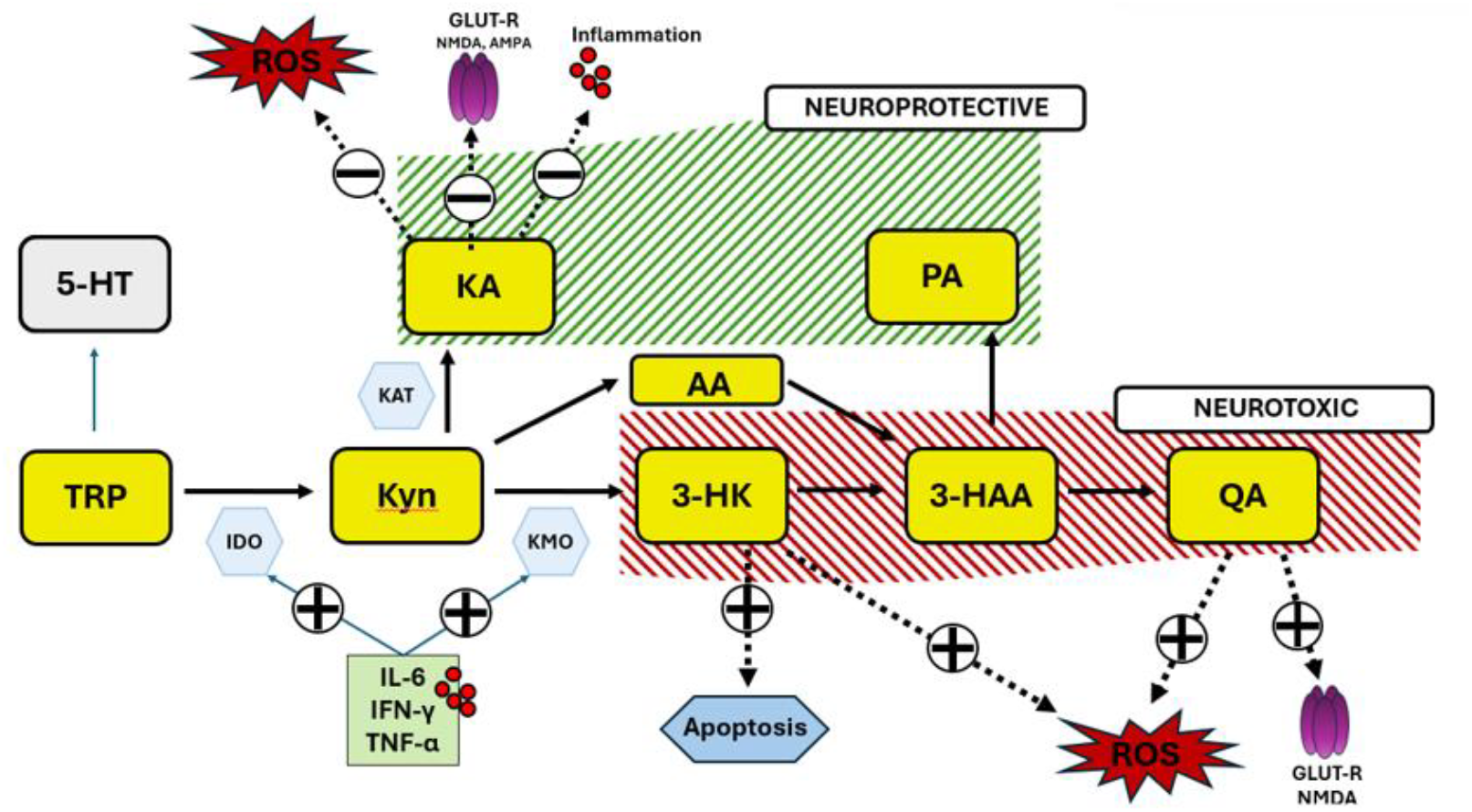
Overview of the Kynurenine Pathway. Metabolites of the Kynurenine Pathway (yellow boxes) divided by the putatively neuroprotective (green) or neurotoxic (red) effects. Tryptophan (**TRP**), Serotonin (**5-HT**), Indoleamine 2, 3 Dioxygenase (**IDO**), Kynurenine (**Kyn**), Kynurenine Aminotransferase (**KAT**), Kynurenic-Acid (**KA**), 3-Hydroxykynurenine (**3-HK**), Kynurenine 3-monooxygenase (**KMO**), Anthranilic Acid (**AA**), Picolinic Acid (**PA**), Quinolinic Acid (**QA**). KA can act as a free radical scavenger and inhibit ROS production, can reduce inflammation (possibly via activity at the Aryl Hydrocarbon Receptor) and act as antagonist of the glutamatergic receptors including the NMDA receptor. In contrast, metabolites of the neurotoxic arm (3-HK, QA) of the pathway can contribute to oxidative stress and QA can act as an NMDA agonist contributing to glutamate induced excitotoxicity.

Few studies [16–18] have examined the relationship between altered KP metabolism and antidepressant response, and none have examined changes in neuroprotective KP metabolite ratios early in the course of treatment or their relationship with brain structure/metabolism. Both hippocampal and amygdala volumes have been associated with response to antidepressant treatment[19–21]. However, longitudinal assessments of KP metabolism and brain structure in the context of SSRI treatment are lacking, despite evidence that peripheral KP metabolism is associated with these volumes[22–25]. Additionally, evidence suggests disrupted glutamatergic signaling is present in MDD (as evidenced by reduced abundance of glutamate [Glu] or a combination of glutamate and glutamine [Glx] in several regions including the anterior cingulate cortex [ACC][26–28]), yet no studies have specifically examined the link between KP metabolites and glutamatergic signaling in the ACC, including in the context of antidepressant treatment. Findings in ketamine studies suggest that NMDA antagonism can increase Glx in specific brain regions[29,30]. While the findings are mixed[31], this raises the possibility that shifting the balance of the KP towards the neuroprotective branch (towards the NMDA antagonist KA) may correct glutamatergic dysfunction and increase Glx in the ACC.

In this longitudinal study, we sought to understand how changes in neuroprotective KP metabolite ratios and activation of the KP were related to response to 8 weeks of SSRI treatment, including early changes (week 4) during the course of treatment, and how changes in KP metabolism would relate to changes in inflammatory markers. Additionally, we sought to understand whether peripheral neuroprotective KP metabolite ratios are related to central measures of brain structure and glutamate metabolism (in the ACC). We hypothesized that response to SSRI treatment would be associated with increases in neuroprotective KP metabolite ratios and that these increases would be associated with increases in hippocampus/amygdala volume as well as increases in Glx in the ACC. We further hypothesized that, in individuals with MDD, higher pre-treatment neuroprotective KP metabolite ratios would be associated with larger amygdalar and hippocampal volumes, as well as greater Glx levels in the anterior cingulate cortex prior to treatment.

## Methods

### Recruitment procedures and study participants

This research was approved by the Institutional Review Board (IRB) of the University of California, San Francisco (UCSF). Participant recruitment was conducted as previously described[32]. Briefly, All MDD subjects were diagnosed with moderate to severe MDD (Hamilton Depression Rating Scale [HDRS] ≥ 17) without psychotic symptoms according to the Structured Clinical Interview for DSM-IV-TR Axis I Disorders (SCID), diagnosis was verified by clinical interview with a board-certified psychiatrist, and participants had no medical or psychiatric co-morbidity. Participants had to pass a urine toxicology screen for drugs of abuse and a urine test for pregnancy for women of child-bearing potential. A subset of these participants has been previously reported on with respect to inflammatory markers (n = 17) [33] and serotonin levels (n = 17) [34] and are presented here in a larger sample for completeness. All KP metabolite data and MR data reported here have not been previously published.

### Measurement of tryptophan, serotonin, and kynurenine metabolites

Kynurenine pathway metabolites, tryptophan, and serotonin were quantified using reverse phase high performance liquid chromatography (HPLC) coupled to a triple quadrupole mass spectrometer (1290 Infinity II LC System, 6470 Triple quadrupole, Agilent Technologies, Santa Clara, CA). Details are included in Supplementary Material.

### MRI Acquisition

MR data were acquired on a 3T Siemens Vision system using a 24-channel transmit-receive head coil (Siemens, Erlangen, Germany). A Magnetization Prepared Rapid Gradient (TR/TE/TI = 2300/2.98/1000 ms, flip angle=9°, voxel resolution=1 mm^3^) sequence was used to acquire 3D sagittal T1-weighted anatomical MR images. These images and those of a turbo spin-echo sequence (TR/TE = 9000/91 ms, flip angle=150°, voxel resolution=0.9 mm x 0.9 mm x 5 mm) were then displayed on the console for the prescription of the volumes-of-interest (VOIs) from which MR spectra were acquired in the same scanning session. Additional details regarding processing of images to obtain amygdala and hippocampal gray matter volumes and details regarding the MR spectroscopy are found in the Supplementary Methods.

### Peripheral Inflammatory Markers

As previously reported, a high sensitivity multiplexed sandwich immunoassay was used to quantify to quantify IL-6 and TNF-α concentrations[33]. CRP was assayed with a latex-enhanced immunoturbidimetric method (Sonora Quest Laboratories). Cytokines were assessed in two different batches. Batch effects were corrected for using the HarmonizR package[35,36].

### Statistical Analysis

Participants with longitudinal pre/post SSRI data was available for 48 individuals. Four participants were missing data for IL-6 and TNF-α and five were missing CRP. Missing data were imputed with the mean and sensitivity analyses without these datapoints showed no impact on results. All data were log transformed prior to analysis. Our *a priori* variables of interest were the KA/3-HK and KA/QA ratios (neuroprotective indices) and Kyn/Trp ratio (reflecting activation of the KP). Responders were defined as ≥ 50% improvement in week 8 HDRS rating relative to baseline and Non-Responders as those with lesser improvement. Delta HDRS (difference between depression severity at week 8 and week 0) was used to examine the relationship between change in kynurenine metabolite ratios and changes in depression severity over the treatment period. Linear mixed effects (LME) model with repeated measures were used to assess for the effect of responder status, time, and the responder x time interaction on the primary outcome measure (Responder Status). A subset of participants (n= 31) had blood assayed at 4 weeks after initiation of treatment and LME model with repeated measures were assessed using all three time points. Models also included age, sex, and BMI as fixed effects covariates and showed no differences from unadjusted models. For clarity, only unadjusted models are presented. To examine changes over time for Responders and Non-Responders separately, post-hoc tests were performed on the estimated marginal means from the LME models using the “emmeans” package in R, with contrasts computed for each group. This approach was also used for the inflammatory markers tested. A subset of the participants had structural MRI data (n = 45) or MRS data (n = 27) at baseline. Relationships between KP metabolite ratios and MR data was assessed via linear regression. Total intracranial volume (ICV) was included as a covariate in MRI sensitivity analyses[25]. Age, sex, and BMI were included as covariates in MRS analyses. Effect sizes for multiple linear regression analyses were calculated via Cohen’s f^2^ statistic[37]. Given *apriori* hypotheses and prior studies linking these kynurenine metabolite ratios to treatment response [16] and brain metrics [25,38,39], we did not correct for multiple comparisons and used a statistical threshold of p<.05 for all tests

## Results

### Participant Characteristics

No differences in age, sex, BMI, pre-treatment depression severity, or antidepressant use were observed between antidepressant Responders and Non-responders. Additional characteristics are detailed in Table 1.

**Table 1.**
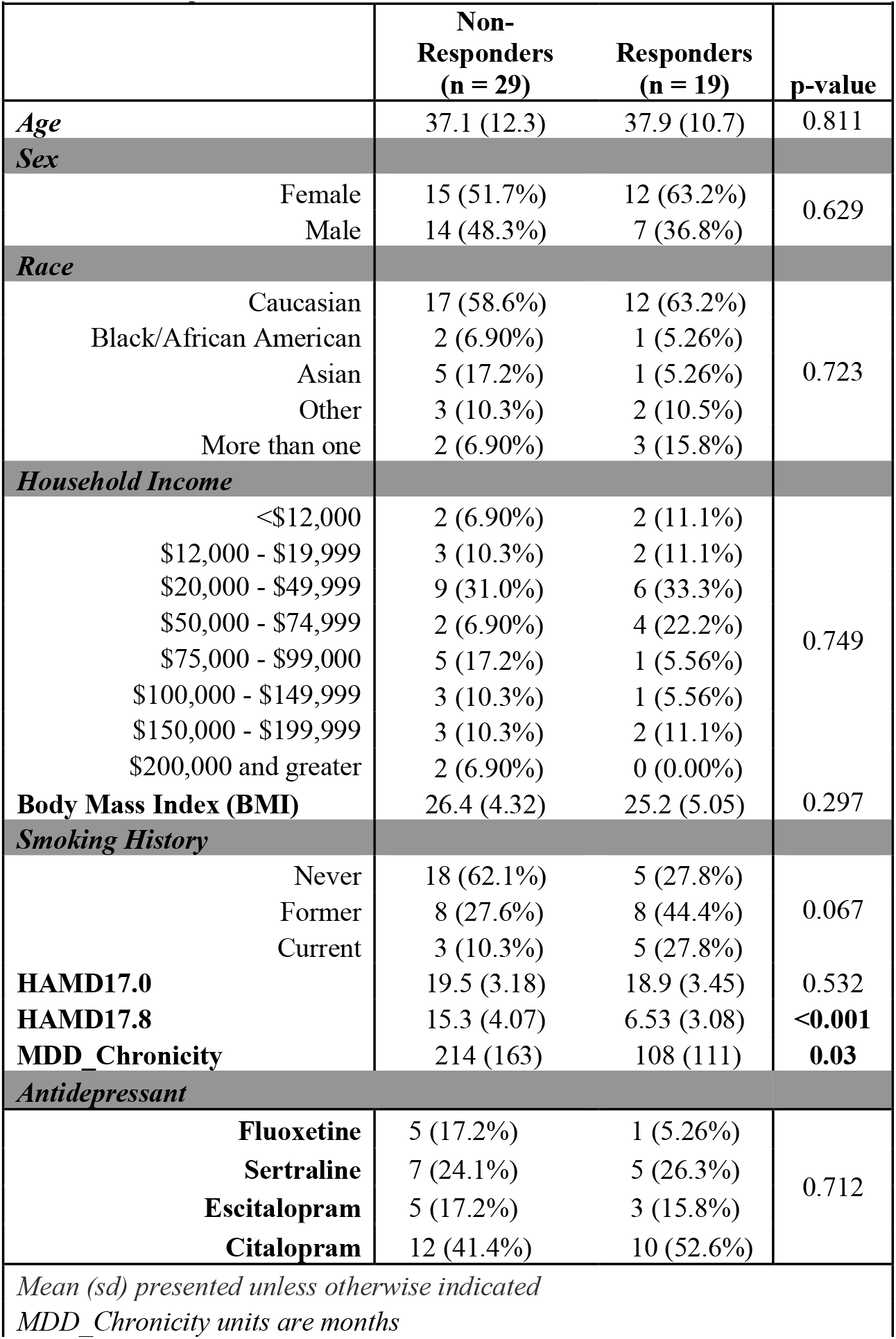
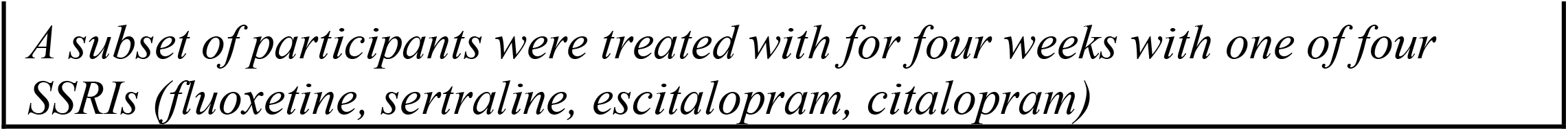
Participant Characteristics.

| Table 1. Participant Characteristics |  |  |  |
| --- | --- | --- | --- |
|  | Non-Responders<br>(n = 29) | Responders<br>(n = 19) | p-value |
| Age | 37.1 (12.3) | 37.9 (10.7) | 0.811 |
| Sex |  |  |  |
| Female | 15 (51.7%) | 12 (63.2%) | 0.629 |
| Male | 14 (48.3%) | 7 (36.8%) |  |
| Race |  |  |  |
| Caucasian | 17 (58.6%) | 12 (63.2%) | 0.723 |
| Black/African American | 2 (6.90%) | 1 (5.26%) |  |
| Asian | 5 (17.2%) | 1 (5.26%) |  |
| Other | 3 (10.3%) | 2 (10.5%) |  |
| More than one | 2 (6.90%) | 3 (15.8%) |  |
| Household Income |  |  |  |
| <\$12,000 | 2 (6.90%) | 2 (11.1%) | 0.749 |
| \$12,000 - \$19,999 | 3 (10.3%) | 2 (11.1%) | |
| \$20,000 - \$49,999 | 9 (31.0%) | 6 (33.3%) | |
| \$50,000 - \$74,999 | 2 (6.90%) | 4 (22.2%) | |
| \$75,000 - \$99,000 | 5 (17.2%) | 1 (5.56%) | |
| \$100,000 - \$149,999 | 3 (10.3%) | 1 (5.56%) | |
| \$150,000 - \$199,999 | 3 (10.3%) | 2 (11.1%) | |
| \$200,000 and greater | 2 (6.90%) | 0 (0.00%) | |
| Body Mass Index (BMI) | 26.4 (4.32) | 25.2 (5.05) | 0.297 |
| Smoking History |  |  |  |
| Never | 18 (62.1%) | 5 (27.8%) | 0.067 |
| Former | 8 (27.6%) | 8 (44.4%) |  |
| Current | 3 (10.3%) | 5 (27.8%) |  |
| HAMD17.0 | 19.5 (3.18) | 18.9 (3.45) | 0.532 |
| HAMD17.8 | 15.3 (4.07) | 6.53 (3.08) | <0.001 |
| MDD_Chronicity | 214 (163) | 108 (111) | 0.03 |
| Antidepressant |  |  |  |
| Fluoxetine | 5 (17.2%) | 1 (5.26%) | 0.712 |
| Sertraline | 7 (24.1%) | 5 (26.3%) |  |
| Escitalopram | 5 (17.2%) | 3 (15.8%) |  |
| Citalopram | 12 (41.4%) | 10 (52.6%) |  |
| Mean (sd) presented unless otherwise indicated |  |  |  |
| MDD Chronicity units are months |  |  |  |

*A subset of participants were treated with for four weeks with one of four SSRIs (fluoxetine, sertraline, escitalopram, citalopram)*

### Response to SSRI treatment is Associated with Increases in Neuroprotective Kynurenine Metabolite Ratios After 8 weeks of Antidepressant Treatment

After 8 weeks of SSRI treatment (n = 48), a significant group x time interaction was observed for the neuroprotective KA/3-HK ratio (*F*_(1,46)_ = 11.92, p = .**001**; **Figure 2A**). Analysis of the effect of time for each group (Responder and Non-Responder) separately showed a significant increase in this ratio in Responders (p = .0006) while Non-Responders showed no difference (p = .362; Post-hoc analyses are detailed in **Supplementary Table 1B**). A similar non-significant pattern was observed for the KA/Kyn ratio (F_(1,46)_ = 3.45 p = *0*.*07;* **Supplementary Table 1A; Supplementary Figure 1A**). Analysis of the effects of time for each group separately showed a significant increase in this ratio in Responders (p = .049) but no significant change in the Non-Responders (p = .650). Serotonin (5-HT) decreased significantly over time for both Responders and Non-responders (F_(1,46)_ = 316.4 p < .*0001*; **Supplementary Table 1A**; **Supplementary Figure 1A**). We also observed that increases in KA/3-HK were associated with decreases in depression severity over the treatment period (β = -0.329 p = .024 and β = -0.86 p = .047; **Supplementary Table 2**) amongst all the depressed participants.

**Figure 2.**
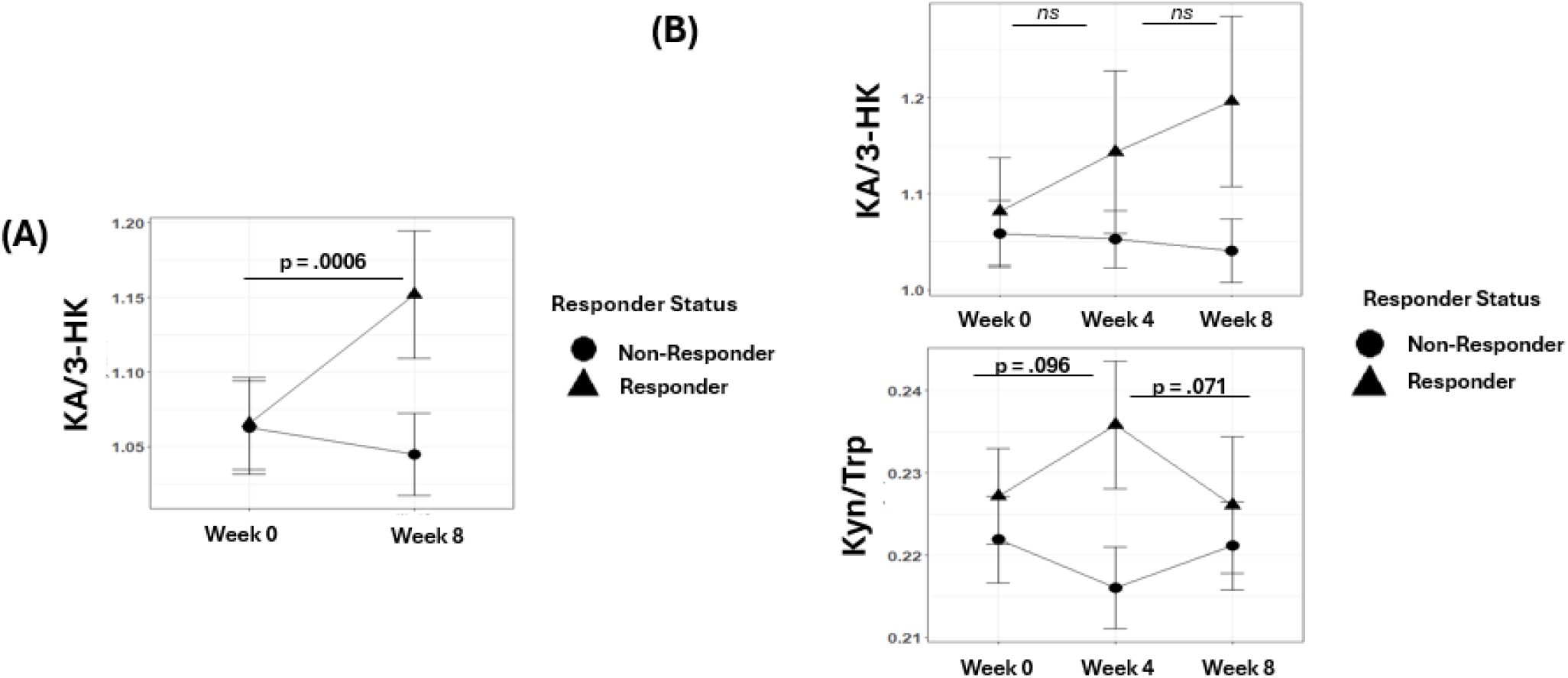
Response to antidepressant treatment is associated with changes in kynurenine metabolism. **(A)** Depressed participants (n = 48) were treated with SSRIs for 8 weeks and response was assessed by change in Hamilton Depression Rating Scale (HDRS). Significant group x time interactions for the KA/3-HK ratio were observed, indicating SSRI responders showed increases in this ratio while Non-responders did not show the same pattern (*F*_*1*,*46*_ *= 11*.*924; p =*.*001*). **(B)** A subset of those treated (n = 31) were assessed 4 weeks. A significant group x time interaction for KA/3-HK was observed, indicating SSRI responders showed an increase by week 8 (*F*_*(2*,*58)*_ *= 4*.*2780 p =* .*018)*. A significant group x time interaction for the Kyn/Trp ratio was observed, indicating SSRI responders had early increases in this ratio which subsequently normalized by week 8 (*F*_*(2*,*58)*_ *= 5*.*2244 p =* .*008*). Post-hoc testing was carried out to identify differences between timepoints for Responders and Non-Responders separately. P-values shown only for Responders as Non-Responders show no significant change over time. For Kyn/Trp ratio, a significant difference was observed at Week 4 between Responders and Non-Responders (t_29_ = -2.5197 p = .018)

### Increased Inflammation is Detected in Non-Responders but not Responders Over 8 Weeks of Treatment

Next, we assessed changes in the abundance of three peripheral inflammatory markers (CRP, IL-6, and TNF-α) over the course of treatment. For IL-6, a significant group x time interaction (F_(1,46)_ = 7.91 p = .007) was observed. Non-Responders showed an increase in IL-6 levels from week 0 to week 8 (p = .0004) while Responders showed no change (**Supplementary Table 3**; **Supplementary Figure 2**). CRP and TNF-α did not change for either Responders or Non-Responders over the treatment period.

### Early Changes in the Kynurenine Pathway is Associated with Treatment Response

A subset of participants with MDD was assessed 4 weeks after initiation of treatment (n = 31; 9 Responders and 22 Non-Responders). A significant group x time interaction was observed for the KA/3-HK ratio (F_(2,58)_ = 4.2780 p = .018; **Supplementary Table 4A**). Among Responders, this ratio had not significantly increased (p = .192) by week 4, but by 8 weeks, it was elevated compared to pre-treatment levels (p = .016; **Figure 2B**; Post-hoc contrasts in **Supplementary Table 4B**). Non-Responders showed no significant changes over the three timepoints.

The Kyn/Trp ratio also displayed a significant group x time interaction (*F*_*(2*,*58)*_ = 5.2244 p = .008; **Figure 2B** and **Supplementary Table 4A**). Responders showed a trend level increase by week 4 (p = .096), followed by a trend level decrease by week 8 (compared to week 4; p= .071; Post-hoc contrasts in **Supplementary Table 4B**). Non-Responders followed a different trajectory. Cross-sectionally, at week 4 Kyn/Trp was higher in Responders compared to Non-Responders (t_9_ = -2.5197 p = .018). A similar group x time interaction was observed for the QA/Trp ratio (F_(2,58)_ = 5.2317 p = .008**; Supplementary Table 4A; Supplementary Figure 1B**). Responders showed a significant increase after 4 weeks (p = .030) followed by a trend level decrease by 8 weeks (p= .063), while Non-Responders showed no significant changes. The 3-HK/Kyn ratio also showed a significant group x time interaction (F_(2,58)_ = 4.764 p = .012; **Supplementary Figure 1B**). In Responders, this ratio increased by week 4 (p = .042) and returned to baseline levels by week 8 (compared to week 4; p = .013) while Non-Responders showed no change. Pre-treatment KP metabolite levels were not different between Responders and Non-Responders (**Supplementary Table 5**).

### Peripheral Neuroprotective Kynurenine Metabolite Ratios are Associated with Amygdalar and Hippocampal Volumes

No changes in amygdalar or hippocampal volumes were observed over the course of 8 weeks and no significant relationships between changes in kynurenine metabolite ratios and changes in amygdala/hippocampal volume were observed over the treatment period (n = 31; **data not shown**). No significant relationships were observed between pre-treatment KA/QA or KA/3-HK and pre-treatment hippocampal volumes (n = 45). However, a significant negative relationship was observed between pre-treatment Kyn/Trp and left hippocampal volume (β =-.320, p = .008, Cohen’s f^2^ = 0.18). Moreover, the neuroprotective KA/QA and KA/3-HK ratios demonstrated positive associations with right amygdalar volume (β = .245, p = .028, Cohens f^2^ = .12; **Table 2**). In our exploratory analysis, we also observed a significant positive relationship between this region and KA/Kyn ratio after adjustment for ICV (β = .317, p = .004). No other kynurenine metabolite ratios were associated with these brain volumes (**Supplementary Table 6**).

**Table 2.**
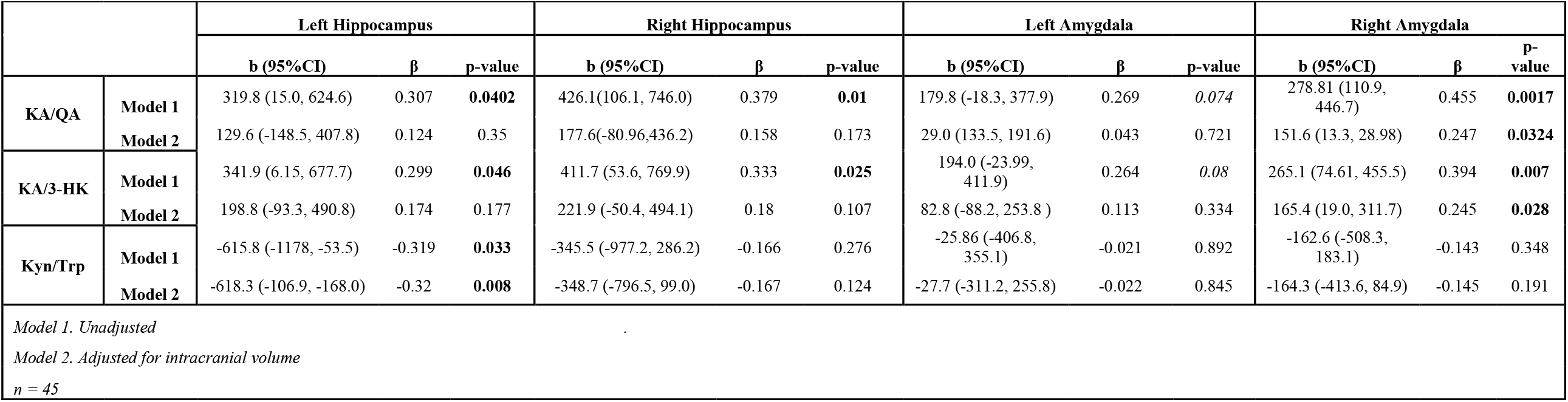
Relationships Between Neuroprotective Kynurenine Metabolite Ratios and Hippocampal/Amygdalar Volumes.

### Peripheral Neuroprotective Kynurenine Metabolite Ratios are Associated with Altered Brain Glutamate Metabolism in the Anterior Cingulate Cortex

We did not observe any changes in Glx in ACC over the course of 8 weeks of antidepressant treatment or any significant relationships between the change in neuroprotective kynurenine metabolite ratios and change in Glx in the ACC (n = 16; **data not shown**). Prior to treatment, we observed a positive relationship between the KA/3-HK ratio and Glx in the ACC (β = .438, p = .022; n = 27), that was further attenuated after adjustment for age, sex, and BMI (β = .307, p = .079, Cohens f^2^ = .15). In addition, a positive relationship was observed between the Kyn/Trp ratio and Glx in the ACC after adjustment for age, sex, and BMI (Unadjusted: β = .043, p = .832; Adjusted: β = .486, p = .02, Cohens f^2^ = .29; **Table 3**). Pre-treatment relationships between additional kynurenine metabolite ratios and ACC Glx are shown in **Supplementary Table 7**. We also examined post-treatment relationships between kynurenine metabolite ratios and ACC Glx in a subset of the participants who were followed longitudinally and had MRS scans (n=16). We observed that the strength of the relationship between the KA/3-HK ratio and Glx in the ACC was increased (β = .668, p = .005) even after adjustment for age, sex, and BMI (β = .652, p = .021) and was associated with a large effect size (Cohens f^2^ = 0.658; **Table 3**). Post-treatment relationships between additional kynurenine metabolite ratios and ACC Glx are detailed in **Supplementary Table 7**. When we restricted the analysis to only those individuals who had assessments at both time points (n = 16), a similar pattern of association was observed. In the pre-treatment samples, no significant relationships were observed between the KA/3-HK ratio and Glx in the ACC and was of a small effect size (β = .321, p = .251; Cohens f^2^ = 0.133; **Supplementary Table 7**). These relationships were noted to be larger in the post-treatment sample (as noted above).

**Table 3.** Relationship between Kynurenine Metabolite Ratios and Glx in the Anterior Cingulate Cortex.

|  |  |  |  |  |
| --- | --- | --- | --- | --- |
|  |  | **b (95%CI)** | **β** | **p-value** |
| **Pre-Treatment** |  |  |  |  |
| **KA/QA** | **Model 1** | 1.02 (-.177, 2.21) | 0.331 | 0.092 |
|  | **Model 2** | .299 (-.961, 1.56) | 0.097 | 0.6275 |
| **KA/3-HK** | **Model 1** | 1.45 (.225, 2.67) | 0.438 | **0.022** |
|  | **Model 2** | 1.01 (-.129, 2.15) | 0.307 | 0.079 |
| **Kyn/Trp** | **Model 1** | .216 (-1.87, 2.30) | 0.043 | 0.832 |
|  | **Model 2** | 2.46 (.433, 4.48) | 0.486 | **0.02** |
| **Post-Treatment** |  |  |  |  |
| **KA/QA** | **Model 1** | 3.48 (.753, 6.20) | 0.591 | **0.016** |
|  | **Model 2** | 3.62 (.707, 6.52) | 0.614 | **0.019** |
| **KA/3-HK** | **Model 1** | 4.46 (1.61, 7.30) | 0.668 | **0.005** |
|  | **Model 2** | 4.35 (.791, 7.91) | 0.652 | **0.021** |
| **Kyn/Trp** | **Model 1** | 3.91 (-2.80, 10.6) | 0.317 | 0.232 |
|  | **Model 2** | 8.78 (2.37, 15.2) | 0.71 | **0.012** |
| *Model 1: Unadjusted*<br>*Model 2: Adjusted for age, sex, BMI*<br>*Pre-treatment data represent a subset of the sample for which MRS data was acquired (n = 27)*<br>*Post-Treatment data represent a subset of the pre-treatment participants treated with an SSRI for 8 weeks (n = 16)* |  |  |  |  |

## Discussion

Our findings demonstrate, for the first time, that response to SSRIs in a medically healthy, unmedicated cohort of individuals with MDD is associated with early activation of the kynurenine pathway, as measured by increased Kyn/Trp ratio at 4 weeks and followed by a shift towards the neuroprotective arm by 8 weeks, extending previous findings from other groups[16]. Lastly, our study demonstrated that greater abundance of neuroprotective metabolites relative to neurotoxic metabolites is associated with larger amygdalar volumes and greater abundance of glutamatergic markers (Glx) in the ACC prior to treatment.

The mechanisms linking SSRI treatment with an increase in the KA/3-HK ratio are not clear but may involve direct effects on Kynurenine Aminotransferase (*KAT*) expression[40], which catalyzes the production of KA from kynurenine. Given the link between inflammation and KP metabolism and the potential anti-inflammatory effects of SSRIs[41], we investigated whether changes in inflammatory markers were associated with changes in neuroprotective kynurenine metabolites. While we found no direct evidence that the increase in KA/3-HK ratio was related to changes in inflammatory markers (data not shown), our assessment of inflammatory markers was limited (IL-6, TNF-α, CRP). Thus, we cannot rule out the possibility that SSRIs induce their effects on markers not assessed here.

We hypothesized that a shift towards the neuroprotective arm of the KP in Responders would be associated with increases hippocampus and amygdala volumes (implicated in MDD pathophysiology and antidepressant response). However, no such changes were observed, possibly due to the study time frame[42] rather than a lack of effects on neurogenesis (as KA can induce BDNF expression[43]). Prior to treatment, neuroprotective indices were associated with larger amygdala volumes while neurotoxic indices and markers of KP activation (Kyn/Trp ratio) were associated with smaller hippocampal volumes, consistent with prior studies[25,38]. Although pre-treatment gray matter volumes were not associated with Responder status in this study (data not shown), larger hippocampal volumes have been linked to superior antidepressant response[44,45]. The amygdala’s role in treatment response is less clear but has been suggested in some studies[46–48]. Our exploratory results are consistent with the hypothesis that increasing flow through the neuroprotective arm of the pathway may facilitate antidepressant response via its effects on neurogenesis [43,49,50], although we cannot address causality. Longitudinal studies over longer time frames are needed to assess the relationship between shifts in KP metabolism and hippocampal/amygdala volumes with treatment.

Given the ability of KP metabolites to modulate NMDA receptor activity and numerous studies demonstrating that MDD is associated with reduced Glx in several brain regions[26], including the ACC[51], we initially hypothesized that successful treatment would be associated with increases in Glx in the ACC. We did not find any evidence of change in Glx over the treatment period (data not shown). However, our results did demonstrate a positive relationship between neuroprotective indices and Glx in the ACC of a medium effect size (Cohen’s f^2^ = .15) in pre-treatment samples. While pre-treatment Glx has not been directly linked to treatment response, ACC activity has been implicated[52–54]. This raises the possibility that the relationship between Glx and KP metabolism in the ACC may influence treatment outcomes.

Interestingly, our results suggest that antidepressant treatment enhances the relationship between KP metabolism and glutamatergic signaling. Notably, in post-treatment samples (compared to pre-treatment) we observed associations with larger effect sizes between neuroprotective KP metabolites and Glx in the ACC. While these analyses utilized all available participants, a sensitivity analysis using only those participants who completed 8 weeks of treatment showed similar results. While statistical tests (data not shown) did not reach conventional significance levels (p <.05), possibly due to our limited sample size, this change in effect size suggests a meaningful change and raises the possibility that treatment enhances the relationship between these systems such that increases in KP metabolism are more tightly linked to changes in glutamatergic signaling markers. This finding warrants future study in larger cohorts to confirm its significance.

Strengths of the study include our relatively medically healthy cohort of unmedicated depressed participants which allowed us to look at the role of depression without significant medical co-morbidity or medication influences. Additionally, our assessment of kynurenine metabolites as well as measures of inflammation allowed us to examine these critical relationships given their role as regulators of KP metabolism. Furthermore, the longitudinal treatment arm of the study (including assessments mid-way through the treatment period) is particularly unique and allowed us to understand how changes in KP metabolism might relate to treatment response including how early changes in these metabolites might relate to treatment response. Our use of peripheral blood measurements coupled with central measures (MRI and MRS) is particularly notable and allows us to draw some parallels between peripheral activity and more central measures.

Limitations of study include the cross-sectional design and modest sample size. Additionally, our assessment of inflammatory markers known to modulate kynurenine metabolism and altered by antidepressants [41,55–58] is limited. Future studies should include comprehensive inflammatory marker panels. Furthermore, we could not directly test the effect of antidepressants on *KAT* enzyme expression and future human studies should examine this relationship. Additionally, we could not measure central kynurenine metabolites and future studies should incorporate these measures to determine links between peripheral and central KP metabolism. Lastly, our use of a composite glutamate and glutamine (Glx) measure limits interpretation of specific glutamate effects. Future studies are needed to disambiguate effects of KP metabolites on glutamate and glutamine using MRS scans (or CSF collection) to estimate ratios of glutamate (and glutamine) to Glx to assess whether disruptions in glutamate neurotransmission are in fact driven by insufficient metabolism of glutamate (or glutamine) in the ACC.

Taken together, these findings raise the possibility that response to SSRIs may arise from shifting the balance of neuroprotective and neurotoxic kynurenine metabolites and highlights putative mechanisms (modulation of brain structure of glutamatergic signaling) by which this balance may contribute to treatment response. Furthermore, these results highlight the possibility that kynurenine metabolism may be useful as an early biomarker of response, facilitating appropriate treatment selection and ultimately more rapid symptom reduction.

## Data/Code Availability

Data/R code available by contacting the authors

## Acknowledgements

The authors acknowledge the assistance of Phuong Hoang, Stacy Ann Miller, the UCSF CTSI Clinical Research Center staff, the PNE Lab research assistants, volunteers and research participants.

## Author Contributions

R.R, K.S., O.W., S.M., V.R., G.W. D.L, L.B., conceived this study. R.R, K.S., O.W, S.M., G.W, analyzed data and generated figures. LB carried out kynurenine assays. R.R., K.S. wrote the manuscript. All authors edited the manuscript.

